# Age-Associated Weaker Immunity to Coronaviruses is Characteristic of Children that Develop Multisystem Inflammatory Syndrome following SARS-CoV-2 Infection

**DOI:** 10.1101/2023.08.28.555120

**Authors:** David Camerini, Antonio Arrieta, Arlo Z. Randall, Johannes S. Gach, Holden Maecker, Janet Hoang, Karen Imfeld, Stephanie Osborne, Claudia Enriquez, Christopher Hung, Joshua Edgar, Adam Shandling, Vu Huynh, Andy A. Teng, Jozelyn V. Pablo, Donald N. Forthal, Joseph J. Campo, Diane Nugent

**Affiliations:** Antigen Discovery Incorporated, Irvine CA; University of California, Irvine, CA; Children’s Hospital of Orange County, Orange, CA; Institute for Immunity, Transplantation, and Infection, Stanford University School of Medicine, Stanford, CA

**Keywords:** MIS-C, SARS-CoV-2, COVID-19, IgG, chemokine, Th1

## Abstract

We analyzed the antibody and cytokine responses of twenty-three patients with multisystem inflammatory syndrome of children (MIS-C) that appeared with a three-to-six-week delay following a mild or asymptomatic SARS-CoV-2 infection. These responses were compared to healthy convalescent pediatric COVID-19 patients approximately twenty-eight days after the onset of symptoms. Both groups had strong IgG responses to SARS-CoV-2 spike (S) and nucleocapsid (N) proteins, but the MIS-C patients had weaker antibody responses to certain epitopes in the SARS-CoV-2 S and N proteins and to the S and N proteins of endemic human coronaviruses (HCoV) compared to pediatric convalescent COVID patients. HCoV antibody reactivity was correlated with age. In contrast, MIS-C patients had elevated serum levels of several proinflammatory cytokines compared to convalescent COVID patients, including interleukins IL-6, IL-8, IL-18 and chemokines CCL2, CCL8, CXCL5, CXCL9 and CXCL10 as well as tumor necrosis factor alpha and interferon gamma. Moreover, many cytokine responses of MIS-C patients were positively correlated with antibody responses to the SARS-CoV-2 S, N, membrane and ORF3a proteins while pediatric convalescent COVID patient cytokine responses were more often negatively correlated with antibody responses to the S, N and ORF3a proteins of SARS-CoV-2.

## Introduction

Since severe acute respiratory syndrome coronavirus type two (SARS-CoV-2) and coronavirus infectious disease of 2019 (COVID-19) were first reported in Wuhan, China, children were recognized to experience milder illness. Later reports suggested that children with underlying conditions, including obesity, were at risk to develop severe disease. Cases of Kawasaki-like illness, frequently complicated by cardiogenic shock, were first reported in children in the United Kingdom in association with SARS-CoV-2 infection several weeks after exposure. Similar cases were reported in the United States (US), and a case definition for multisystem inflammatory syndrome in children (MIS-C) was developed. This unique and severe disease usually appears three to six weeks following a typically asymptomatic SARS-CoV-2 infection and is characterized by a variety of symptoms and signs including fever, pain, rash, vomiting and diarrhea. Cardiovascular involvement, often with shock, is the most common severe manifestation of MIS-C, followed by respiratory disease, coagulopathy, hematological abnormalities (mainly thrombocytopenia), gastrointestinal disease, acute kidney injury and neurologic involvement (Miller et al., 2022)

The US Centers for Disease Control and Prevention recently reported that there were 4470 cases of MIS-C in the USA from February 2020 to July 2021 (Miller et al., 2022). The average age of these MIS-C patients was 9 years and almost 60% were male. The incidence of MIS-C per 100,000 children was 54.5 during the Alpha wave, 49.2 amid Delta, and 3.8 in the Omicron era, despite the fact that the majority of children remained unvaccinated. Compared with the Omicron period, the incidence of MIS-C was 14.3 times higher amid Alpha and 12.9 times higher during Delta. As the incidence of MIS-C has declined during the pandemic with the evolution of viral variants, it is likely that changes in viral antigens and/or lack of prior immunity resulted in triggering the sustained inflammatory response seen in MIS-C (Holm et al., 2022; Levy et al., 2022).

Herein, we compared antibody responses to key viral antigens in patients with MIS-C and in convalescent pediatric patients following acute symptomatic COVID-19. We also examined cytokine responses in the two groups of patients and the correlations of cytokine responses to antibody responses to viral antigens. Our findings suggest that there are differences in both antibody and cytokine responses to SARS-CoV-2 infection in patients with MIS-C and following acute disease.

## Materials and Methods

### Patients

Children admitted with evidence of recent SARS-CoV-2 infection who met CDC criteria for MIS-C were approached for participation in our SARS-CoV-2/MIS-C study. Control patients were part of an ongoing prospective, open-label COVID convalescent plasma (CCP) treatment trial being conducted at Children’s Hospital of Orange County (CHOC), Orange, California. We obtained approval under eINDs through the FDA as well as from our local institutional review board. Patients≤26 years old, with COVID-19 confirmed via positive SARS-CoV-2 polymerase chain reaction (PCR) test in nasopharyngeal (NP) swab were eligible if they met any of the following: (1) hospitalized requiring oxygen; (2) oxygen saturation≤93% on ambient air; (3) partial pressure aO2:FIO2 ratio <300; and/or (4) pulmonary infiltrates >50% within 24-48 hours of admission. Signed informed consent (and assent when indicated) was obtained from subjects that agreed to participate in the antibody kinetics study; receiving CCP did not require informed consent. Subjects received CCP at 10ml/kg up to 1 unit (approximately 260 ml. Demographic data and co-morbidities were abstracted from the hospital’s electronic medical records. Comorbidities were identified by documented primary diagnosis with associated ICD10 codes and/or health encounters at related clinics or by record of visits to specialty care clinics for treatment of asthma, diabetes, neurologic, oncologic, cardiac, or hematologic conditions; obesity was defined as BMI ≥95^th^ percentile for age-gender^22-24^. Per American Red Cross Guidelines, donors were eligible to provide CCP if: (a) they were initially proven positive for SARS-CoV-2 by a laboratory test; and either (b1) ≥ 14 days from symptom resolution with repeat documented negative test for SARS-CoV-2, or (b2) ≥ 28 days from symptom resolution without repeat test results at the time of plasma collection.

### Serum/Plasma samples

Samples were obtained from MIS-C patients on admission and from convalescent pediatric COVID patients 21 days post infusion which was an average of 28 days post onset of symptoms. Blood samples were collected, serum was separated and kept frozen at -80oC until processed.

### Pseudotyped virus production and Virus neutralization assay

SARS-CoV-2 pseudotyped lentiviral vector virions were generated by co-transfecting 293-T cells with *HIV-1*_*NL4-3/*_*gag-iGFP*_*/ΔEnv*_ plasmid as well as pcDNA 3.1 SARS CoV-2 S. Plasmids were mixed with polyethylenimine (PEI) at a DNA/PEI ratio of 1:3 an added to the cells. After 72-96 hours cell supernatants were harvested and cleared from cells. Virus aliquots were stored at -80°C. Multi-well flat bottom tissue culture plates were used to mix heat inactivated and diluted patient plasma samples with SARS-CoV-2 pseudotyped lentiviral vector virions. After 1 hour at 37°C, ACE2 expressing HEK 293T cells (BEI Resources), were added and plates were incubated for 72 hours. Cells were then detached, washed once with DPBS, and fixed with 4% paraformaldehyde. Target cells were subsequently analyzed by flow cytometry for green fluorescent protein expression. The median fluorescence intensity of infected cells without serum was used to determine the ID_50_ of serum samples.

### Virus capture assay

Capture assays were performed as previously described (Gach et al., 2017). In brief, ELISA plates were coated overnight with rabbit anti human Fc gamma chain-specific antibody (Jackson ImmunoResearch). Plates were then washed with DPBS, blocked with 5% nonfat dry milk in DPBS, and incubated for 1 hour at 37°C. After plates were washed, 50 μL of diluted patient plasma samples (1:50 in DMEM medium containing 10% FBS) were added. One hour later unbound antibodies were removed and SARS-CoV-2 pseudotyped HIV-1 virions were added for 4 hours at 37°C. After washing five times with DPBS, bound virions were lysed with lysis buffer (ZeptoMetrix) and incubated for 5 min at room temperature. Lysed samples were then transferred into storage plates and kept at -20°C until p24 analysis. The HIV-1 p24 content of the captured vector virions was determined using the Retro-Tek HIV-1 p24 antigen ELISA kit (ZeptoMetrix) following the manufacturer’s instructions. Levels of p24 were calculated by a point-to-point algorithm.

### SARS-CoV-2 Surrogate Virus Neutralization Test (sVNT)

The SARS-CoV-2 surrogate virus neutralization test (GenScript) was used to detect neutralizing antibodies targeting the viral spike (S) protein receptor binding domain. The test is based on antibody-mediated blockage of the interaction between the angiotensin converting enzyme 2 (ACE2) receptor protein and the receptor binding domain. The test was performed according to manufacturer’s instructions. Samples and controls were tested in duplicate. Cutoff values were determined according to manufacturer’s instructions. The percent inhibition was calculated as [1− (OD of sample/OD of negative control)] x 100.

### Protein microarray analysis of serum samples

The multi-coronavirus protein microarray, produced by Antigen Discovery, Inc. (ADI, Irvine, CA, USA), included almost one-thousand features including full-length coronavirus proteins, overlapping 100, 50 and 30 aa protein fragments and overlapping 13-20 aa peptides from SARS-CoV-2 (WA-1), SARS-CoV, MERS-CoV, HCoV-NL63 and HCoV-OC43. Purified proteins and peptides were obtained from BEI Resources. All these coronavirus proteins were produced in *Escherichia coli* except the SARS-CoV-2 and SARS-CoV S proteins, which were made in Sf9 insect cells and the SARS-CoV-2 RBD, made in HEK-293 cells. Other proteins and protein fragments were expressed using an *E. coli in vitro* transcription and translation (IVTT) system (Rapid Translation System, Biotechrabbit, Berlin, Germany) and printed on nitrocellulose-coated glass AVID slides (Grace Bio-Labs, Inc., Bend, OR, USA) using an Omni Grid Accent robotic microarray printer (Digilabs, Inc., Marlborough, MA, USA). Microarrays were probed with sera and antibody binding detected by incubation with fluorochrome-conjugated goat anti-human IgG (Jackson ImmunoResearch, West Grove, PA, USA or Bethyl Laboratories, Inc., Montgomery, TX, USA). Slides were scanned on a GenePix 4300A High-Resolution Microarray Scanner (Molecular Devices, Sunnyvale, CA, USA), and raw spot and local background fluorescence intensities, spot annotations and sample phenotypes were imported and merged in R (R Core Team, 2017), in which all subsequent procedures were performed. Foreground spot intensities were adjusted by subtraction of local background, and negative values were converted to one. All foreground values were transformed using the base two logarithm. The dataset was normalized to remove systematic effects by subtracting the median signal intensity of the IVTT controls for each sample. With the normalized data, a value of 0.0 means that the intensity is no different than the background, and a value of 1.0 indicates a doubling with respect to background. For full-length purified recombinant proteins and peptide libraries, the raw signal intensity data was transformed using the base two logarithm for analysis.

### Serum Cytokine Quantitation

Cytokine levels in serum samples were measured by multiplex Luminex assay at the Stanford Human Immune Monitoring Core laboratory using established protocols. Kits were purchased from EMD Millipore Corporation, Burlington, MA., and run according to the manufacturer’s recommendations with modifications described as follows: H80 kits include 3 panels: Panel 1 is Milliplex HCYTA-60K-PX48. Panel 2 is Milliplex HCP2MAG-62K-PX23. Panel 3 includes the Milliplex HSP1MAG-63K-06 and HADCYMAG-61K-03 (Resistin, Leptin and HGF) to generate a 9 plex. The assay setup followed recommended protocol: Briefly: samples were diluted 3-fold (Panel 1&2) and 10-fold for Panel 3. 25ul of the diluted sample was mixed with antibody-linked magnetic beads in a 96-well plate and incubated overnight at 4°C with shaking. Cold and Room temperature incubation steps were performed on an orbital shaker at 500-600 rpm. Plates were washed twice with wash buffer in a BioTek ELx405 washer (BioTek Instruments, Winooski, VT). Following one-hour incubation at room temperature with biotinylated detection antibody, streptavidin-PE was added for 30 minutes with shaking. Plates were washed as described above and PBS added to wells for reading in the Luminex FlexMap3D Instrument with a lower bound of 50 beads per sample per cytokine. Each sample was measured in duplicate wells. Custom Assay Chex control beads were purchased and added to all wells (Radix BioSolutions, Georgetown, Texas). Wells with a bead count <50 were flagged, and data with a bead count <20 were excluded.

### Statistical Analysis

Means and 95% confidence intervals were calculated from patient clinical lab test results. Student’s t-tests were used for comparison of the individual antibody response means between groups. Correlation of protein microarray and cytokine ELISA measurements used Pearson’s correlation coefficient (*ρ*). Adjustment for the false discovery rate was performed using the “p.adjust” function in R (Benjamini and Hochberg, 1995). Data visualization was performed using the ComplexHeatmap (Gu et al., 2016), ggplot2 and the heatmap.2 function of gplots (Warnes et al., 2022) packages in R. Unadjusted *p*-values (P_raw_) are shown in graphics, and significance of *p*-values that were adjusted for multiple comparisons is shown in blue asterisks. Multivariable linear regression models were fit to antibody data with age in years as a covariate, due to significant associations with age in stratified univariate models and fit to cytokine data with age and BMI as covariates, due to the association of Leptin with BMI. For figure 2, *p*-values were calculated using a nonparametric Mann-Whitney ranks test (GraphPad Prism 9.2).

## Results

### Clinical course and clinical lab results

We identified and enrolled 23 MIS-C patients with evidence of recent SARS-CoV-2 infection who met CDC criteria for MIS-C and 10 previously healthy, immune competent pediatric patients who had recovered from symptomatic COVID-19 following hospitalization. Both groups received treatment at Children’s Hospital of Orange County (CHOC) in Orange, CA, USA. MIS-C patients had their blood drawn before receiving treatment with intravenous immunoglobulin, solumedrol and enoxaparin. Convalescent COVID patients were previously treated with remdesivir, dexamethasone, convalescent plasma (CCP) and/or a combination of these therapies during acute disease and followed for three months following therapy. Longitudinal analysis of IgG reactivity in patients that received CCP indicated that antibodies from infusion constituted a small fraction of the SARS-CoV-2 reactive IgG on day 28 post onset of symptoms (Fig. S1). Consistent with previous reports (Miller 2021), the MIS-C patients we studied were significantly younger than the pediatric COVID patients we treated. In part as a result of this age difference, the MIS-C patients also had significantly lower average body mass index (BMI) scores. Moreover, compared to convalescent pediatric COVID patients on day 28 post onset of symptoms, MIS-C patients had significantly fewer platelets and elevated C-reactive protein (CRP) on admission to hospital (Table).

**Table.**
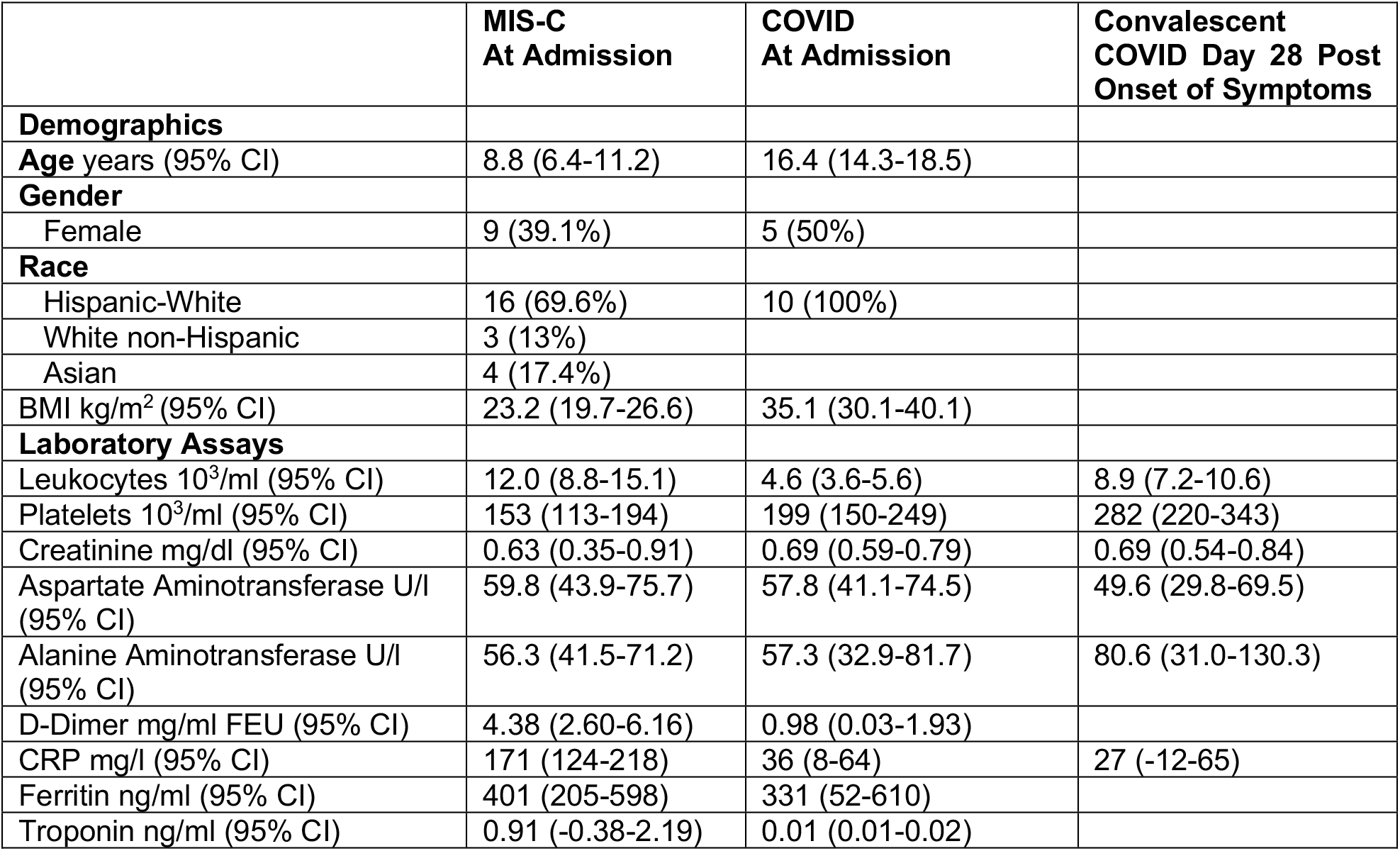
Patient demographics and mean clinical lab values.

**Table. Patient demographics and mean clinical lab values**
|  | **MIS-C**<br>**At Admission** | **COVID**<br>**At Admission** | **Convalescent**<br>**COVID Day 28 Post**<br>**Onset of Symptoms** |
| --- | --- | --- | --- |
| **Demographics** |  |  |  |
| **Age** years (95% CI) | 8.8 (6.4-11.2) | 16.4 (14.3-18.5) |  |
| **Gender** |  |  |  |
| Female | 9 (39.1%) | 5 (50%) |  |
| **Race** |  |  |  |
| Hispanic-White | 16 (69.6%) | 10 (100%) |  |
| White non-Hispanic | 3 (13%) |  |  |
| Asian | 4 (17.4%) |  |  |
| **BMI** kg/m<sup>2</sup> (95% CI) | 23.2 (19.7-26.6) | 35.1 (30.1-40.1) |  |
| **Laboratory Assays** |  |  |  |
| Leukocytes 10<sup>3</sup>/ml (95% CI) | 12.0 (8.8-15.1) | 4.6 (3.6-5.6) | 8.9 (7.2-10.6) |
| Platelets 10<sup>3</sup>/ml (95% CI) | 153 (113-194) | 199 (150-249) | 282 (220-343) |
| Creatinine mg/dl (95% CI) | 0.63 (0.35-0.91) | 0.69 (0.59-0.79) | 0.69 (0.54-0.84) |
| Aspartate Aminotransferase U/l (95% CI) | 59.8 (43.9-75.7) | 57.8 (41.1-74.5) | 49.6 (29.8-69.5) |
| Alanine Aminotransferase U/l (95% CI) | 56.3 (41.5-71.2) | 57.3 (32.9-81.7) | 80.6 (31.0-130.3) |
| D-Dimer mg/ml FEU (95% CI) | 4.38 (2.60-6.16) | 0.98 (0.03-1.93) |  |
| CRP mg/l (95% CI) | 171 (124-218) | 36 (8-64) | 27 (-12-65) |
| Ferritin ng/ml (95% CI) | 401 (205-598) | 331 (52-610) |  |
| Troponin ng/ml (95% CI) | 0.91 (-0.38-2.19) | 0.01 (0.01-0.02) |  |

### Patient antibody reactivity with coronavirus antigens

MIS-C patients had significantly less IgG reactive with the SARS-CoV-2 S, S2 and S2’ proteins and several SARS-CoV-2 N and S protein fragments as well as HCoV N and S2 proteins (Fig. 1A). These included several carboxy-terminal fragments of the SARS-CoV-2 S protein, within amino acid (aa) 501 to aa 588 of the membrane proximal connector region of the S2 domain, and several fragments of the N protein, in the dimerization domain, aa 201-300 and C-terminal domain aa 301-419 (Peng et al., 2020). Moreover, MIS-C patient IgG was significantly less reactive with full-length HCoV-229E and HCoV-NL63 N proteins and with HCoV-OC43 S2 protein.

**Figure 1.**
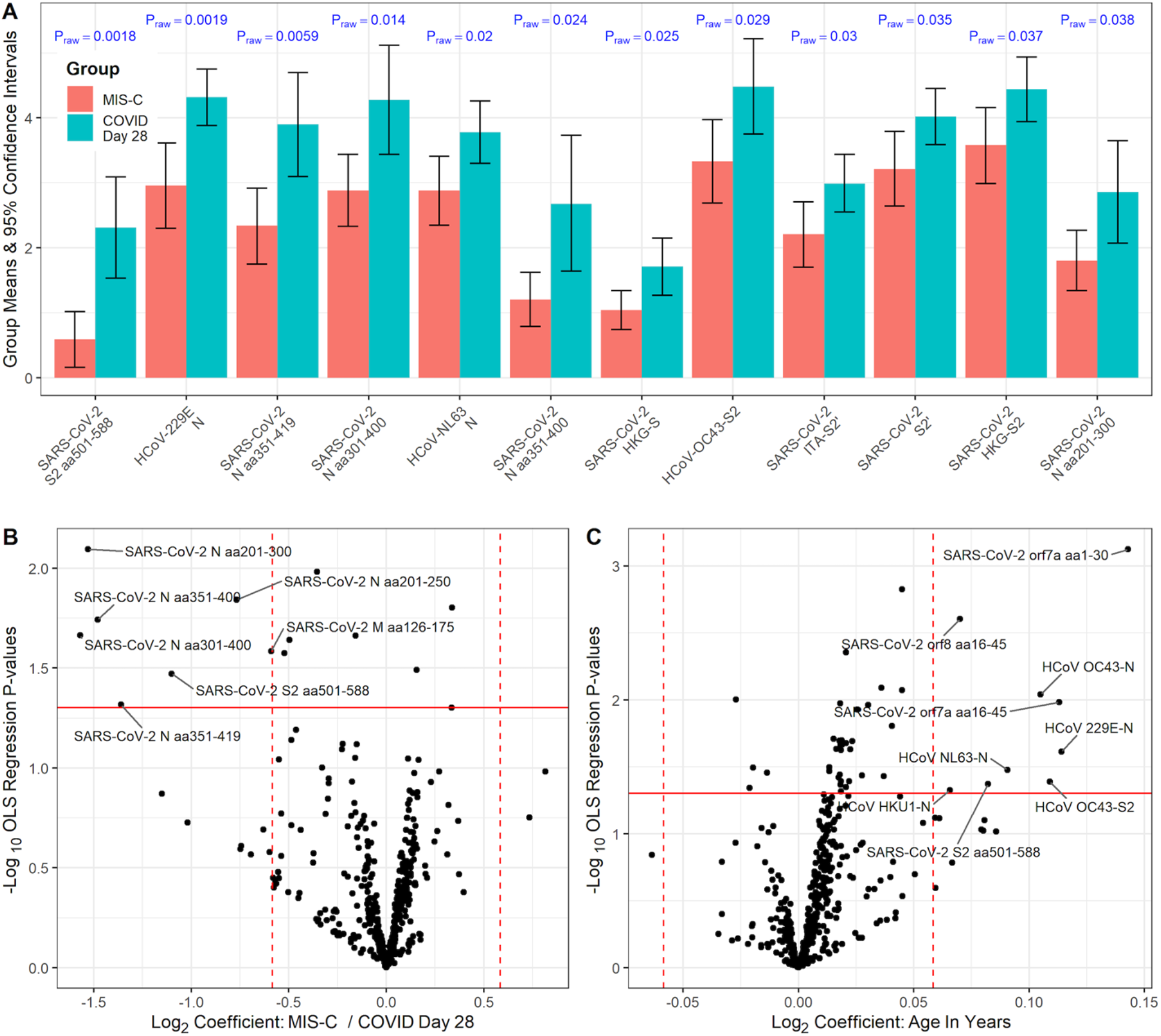
MIS-C patient serum IgG was significantly less reactive with several coronavirus antigens than day 28 convalescent COVID sera. (A) The top twelve most differentially reactive antigens among MIS-C patient sera compared to convalescent pediatric COVID patient sera on day 28 post onset of symptoms are shown. They are ranked from left to right based on their t-test p values. Normalized group means and 95% confidence intervals are shown on a base 2 log scale. (B-C) The effect of MIS-C vs. COVID convalescence and age on antibody levels using multivariable ordinary least squares (OLS) linear regression is shown in the volcano plots for 463 full length or fragmented coronavirus proteins. The horizontal red lines represent an inverse log_10_ OLS regression P-value of 0.05 (values above are P<0.05). The vertical dashed red lines in B represent 50% difference in antibody levels between groups and in C a 50% change in antibody levels per 10 years of age. Antigens with P-values < 0.05 and outside of the vertical lines are labeled. (B) the x-axis represents the effect estimate of MIS-C patient sera vs. day 28 convalescent pediatric COVID patient sera adjusted for age; values less than zero indicate greater antibody reactivity in convalescent COVID day 28 patient sera. (C) the x-axis represents the effect of each year of age on antibody levels in both patient groups.

**Figure 2.**
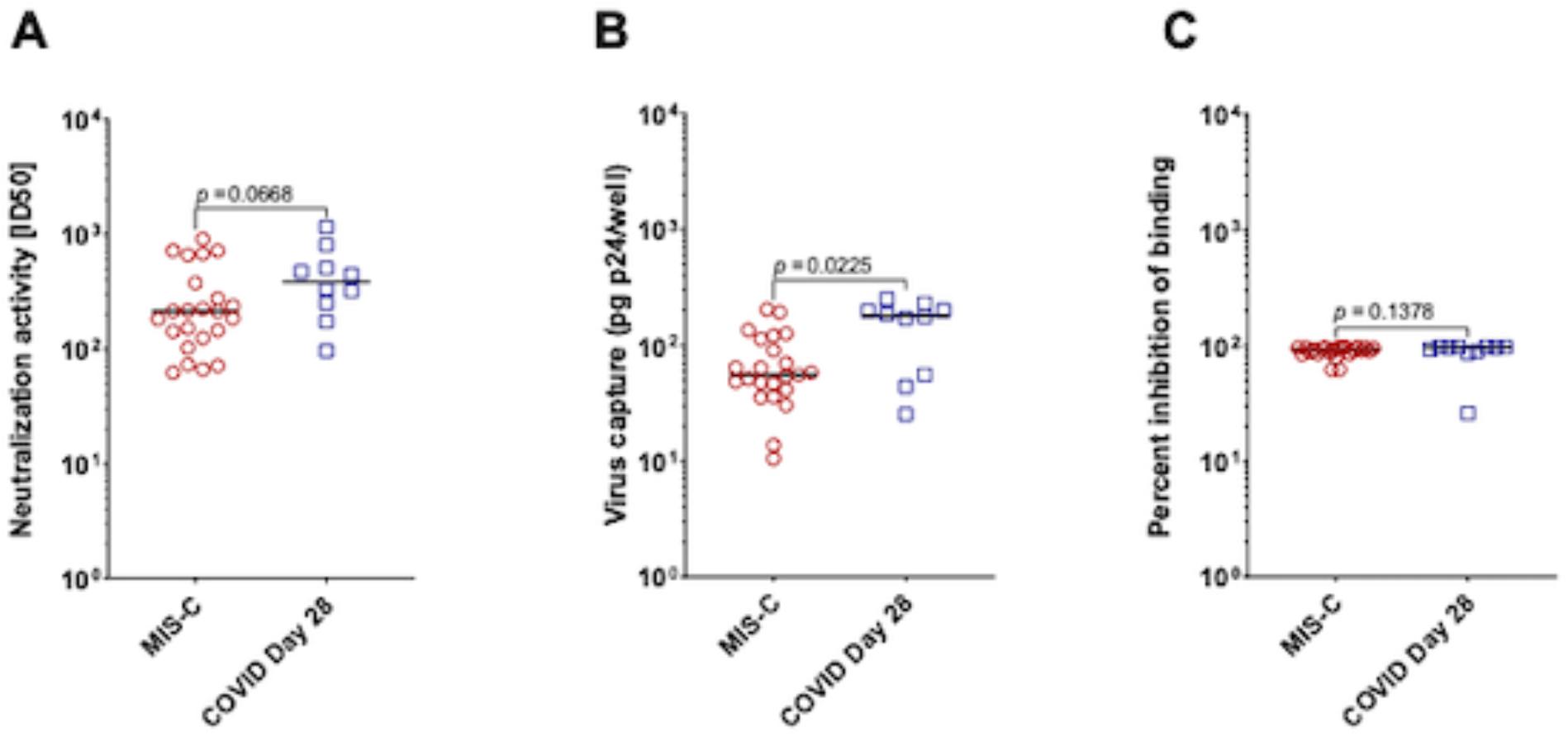
MIS-C patient sera exhibit similar SARS CoV-2 neutralization, but less virion capture activity compared to convalescent pediatric COVID-19 sera. (A) No significant difference was found between the median ID_50_ activity of MIS-C patients and the pediatric control COVID-19 patients. (B) Pediatric control COVID-19 patients captured significantly more p24 than the MIS-C patients. (C) No difference was seen in the ability of the two groups of sera to block binding of SARS-CoV-2 S protein RBD to ACE2. P-values were calculated using a nonparametric Mann-Whitney ranks test (GraphPad Prism 9.2). Lines represent median ID_50_’s, bound p24 and percent inhibition respectively.

We used multivariate models to examine the effects of age, sex and BMI on patient antibody reactivity with coronavirus antigens; only age had significant effects. When the effects of age were excluded, only the fragments of the SARS-CoV-2 N and S2 proteins listed above and one M protein fragment, aa 126-175 in the beta-sheet domain (Zhang et al., 2022), were significantly differentially reactive with MIS-C patient IgG compared to convalescent pediatric COVID-19 patient IgG (Fig. 1B). These regions are not sites of diversity between SARS-CoV-2 Wuhan 1 and the Omicron lineage variant BA.5 except for three amino acid substitutions in the N protein; R203K and G204R in the dimerization domain as well as S413R near the C-terminus. Patient IgG reactivity with HCoV N and S2 proteins as well as several SARS-CoV-2 S2 and accessory protein fragments was significantly affected by age (Fig. 1C). Greater than half of the antibody responses against reactive antigens on the array were normally distributed by Shapiro-Wilk test (Fig. S2). Analysis using nonparametric statistical methods yielded similar results to the parametric methods presented (Fig. S3). These data suggest that lower levels of preexisting antibodies to HCoV’s and SARS-CoV-2 proteins in younger children may play a role in the etiology of MIS-C following SARS-CoV-2 infection.

### SARS-CoV-2 Neutralizing Activity of Patient Sera

Both MIS-C sera and convalescent pediatric COVID sera exhibited substantial SARS-CoV-2 neutralizing activity, and there was no significant difference between the neutralizing activity of the two groups (Fig. 2A). Conversely, the median SARS-CoV-2 pseudovirus capture activity was significantly more prominent in the convalescent pediatric COVID-19 patient sera (Fig. 2B). This likely reflects antibody binding to regions of the SARS-CoV-2 S protein other than the receptor binding domain (RBD), since both groups of sera blocked binding of the RBD to ACE2 equally well (Fig. 2C).

### Patient cytokine responses

MIS-C patients had a significantly different repertoire of cytokines in their serum compared to convalescent pediatric COVID-19 patients. Four cytokines were at least four-fold higher in MIS-C subjects than in convalescent COVID-19 sera on day 28: CXCL9, CXCL10, IL-10, and IL-6 (Fig. 3). Three of these, CXCL9, CXCL10 and IL-6 are pro-inflammatory and characteristic of Th1 responses, while IL-10, paradoxically is anti-inflammatory for Th1 immune responses. Most of the other cytokines that are significantly elevated in MIS-C patients are pro-inflammatory, however, including IL-18, CCL2, TNF-α and CCL8 (Dinarello, 2000). This is consistent with the systemic inflammation that characterizes the clinical course of MIS-C. Four cytokines were at least four-fold higher in COVID sera day 28 than MIS-C: CCL22, CCL17, Leptin and CXCL5. We used a multivariable model to assess the effect of age and BMI on cytokine secretion and found few significant effects (Fig S4A). There were no significant effects of age alone (Fig S4B), however, the higher leptin signal in the convalescent COVID patients is associated with the higher BMI of these patients (Fig. S4C). Many other statistically significant differences in cytokine levels between MIS-C patients and convalescent pediatric COVID patients were also found (Fig. 3). As with the antibody analysis, approximately half of the cytokines tested were normally distributed (Fig. S5), and nonparametric statistical methods yielded similar results (Fig. S6).

**Figure 3.**
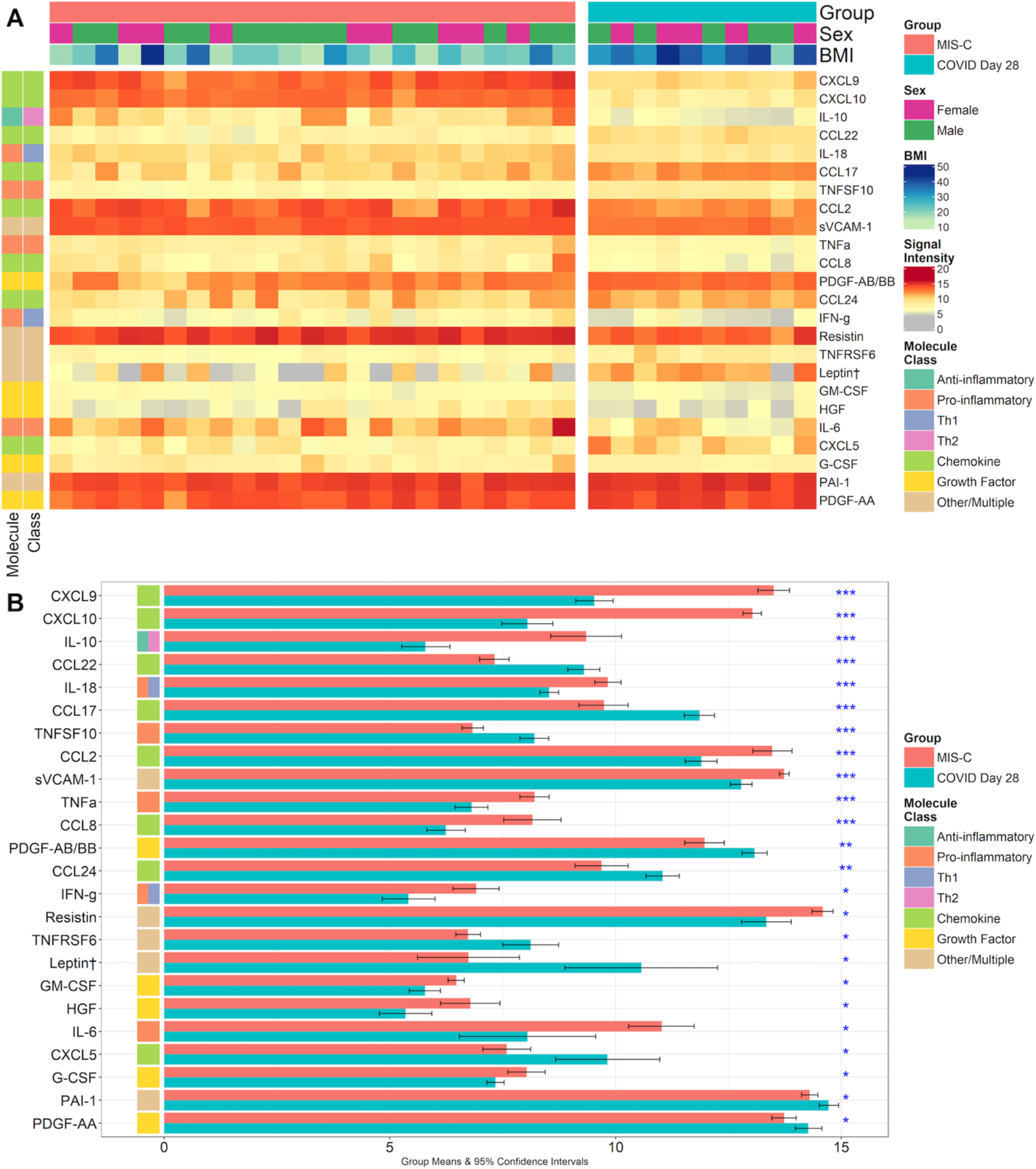
Cytokines with the most significant mean differences between the MIS-C samples and convalescent pediatric COVID samples. (A) The heatmap shows the twenty cytokine signals that were most significantly different between the two groups with significance decreasing from top to bottom. The color key denotes the signal intensity on a log_2_ scale. Columns in each patient group display data from a single patient. (B) The bar graph shows the group means and 95% confidence intervals for each of the twenty-four most significantly differentially expressed cytokines in the group comparison on a log_2_ scale. The significance of the difference between the group means was adjusted for multiple comparisons and is indicated by asterisks; *<0.05, **<0.005 and ***<0.0005. † Leptin was significantly associated with BMI and not significantly different between the two patient groups after adjustment (Fig. S4).

### Correlation of patient antibody and cytokine responses

MIS-C patient IgG binding to SARS-CoV-2 S1 protein was significantly positively correlated with Th1-biased proinflammatory cytokine and chemokine secretion, particularly with IL-6, CCL8, IFN-γ and TNF-α (Fig. 4). In contrast, convalescent COVID patient IgG binding to SARS-CoV-2 proteins S2, S2’ and N was negatively correlated with the levels of many cytokines and positively correlated only with CCL22. This pattern of correlations between antibody binding and cytokine levels, mostly positive for MIS-C patients and predominantly negative for convalescent COVID patients, continued when we analyzed antibody binding to reactive fragments of the SARS-CoV-2 S, N, M, ORF3a, ORF7 and ORF8 proteins (Fig. 5). MIS-C patient IgG binding to SARS-CoV-2 S and S1 proteins and to fragments in the carboxy (C) terminal region of the S1 protein from aa 500 to 600 and in the C-terminal region of the S2 protein, aa 400 to 588, was strongly positively correlated with levels of IL-6, CCL8, TNF-α, IFN-γ, IL-18, soluble ICAM-1, CXCL8 and CCL2. MIS-C patient IgG binding to SARS-CoV-2 nucleocapsid protein fragments, covering amino acids 326 to 375 and 376 to 419 was, however, negatively correlated with levels of several cytokines including CCL26, FGF-β, IL-1β, IL-12, LIF and CCL3. Convalescent pediatric COVID patient IgG binding to many fragments of the SARS-CoV2 S2 protein was negatively correlated with levels of several cytokines, most significantly G-CSF, CCL5, IL-1α and CXCL13.

**Figure 4.**
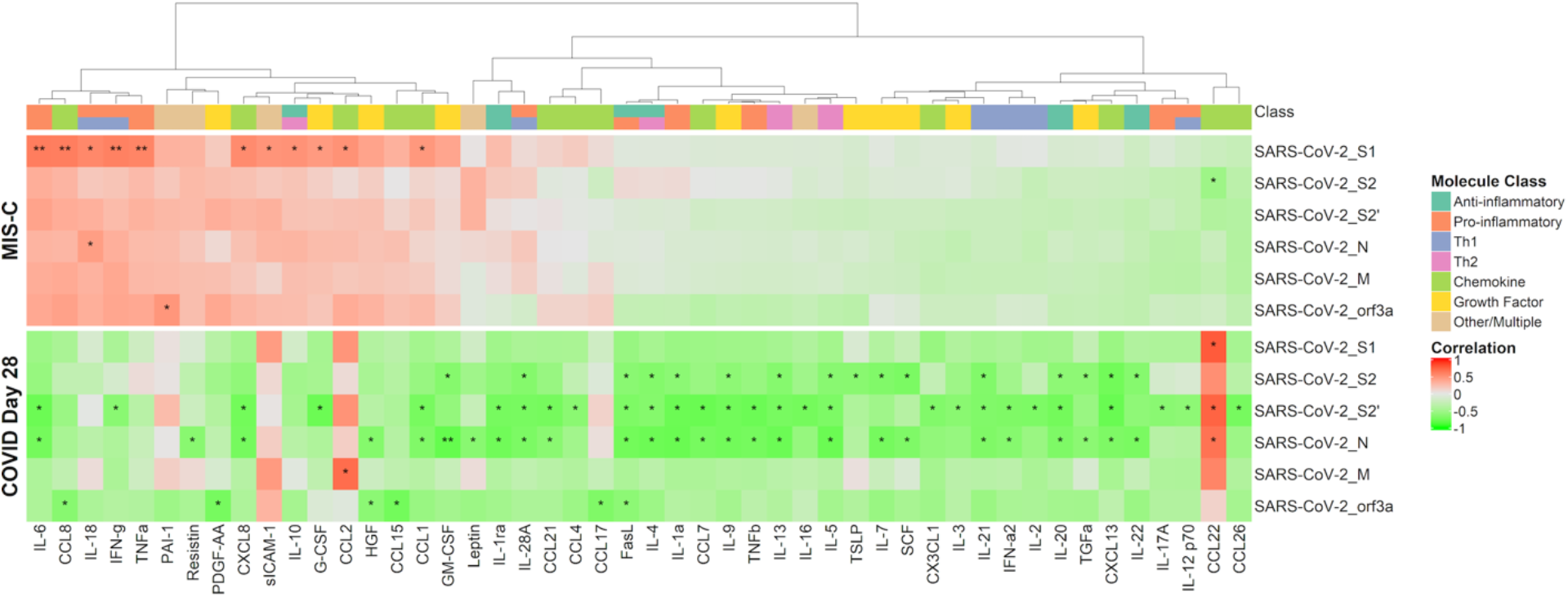
Correlation between IgG binding to SARS-CoV-2 proteins with selected cytokine levels. The heatmap shows the Pearson’s correlation coefficient between antibody and cytokine levels on a colorimetric scale. Significance of the correlations are shown by asterisks (*p<0.05, **p<0.005). Plots are separated by patient group; MIS-C (top) and convalescent COVID-19 at 28 days post-onset of symptoms (bottom). Antigens displayed correspond to full-length proteins produced *in vitro* that were seropositive (≥1.0 normalized log_2_ signal intensity) in at least 20% of the study population. Cytokines were included if at least one significant association with the selected antibody responses was observed in either group of children. Cytokine classes are annotated at the top of the heatmap. The dendrogram at the top of the heatmap represents hierarchical clustering on the MIS-C group.

**Figure 5.**
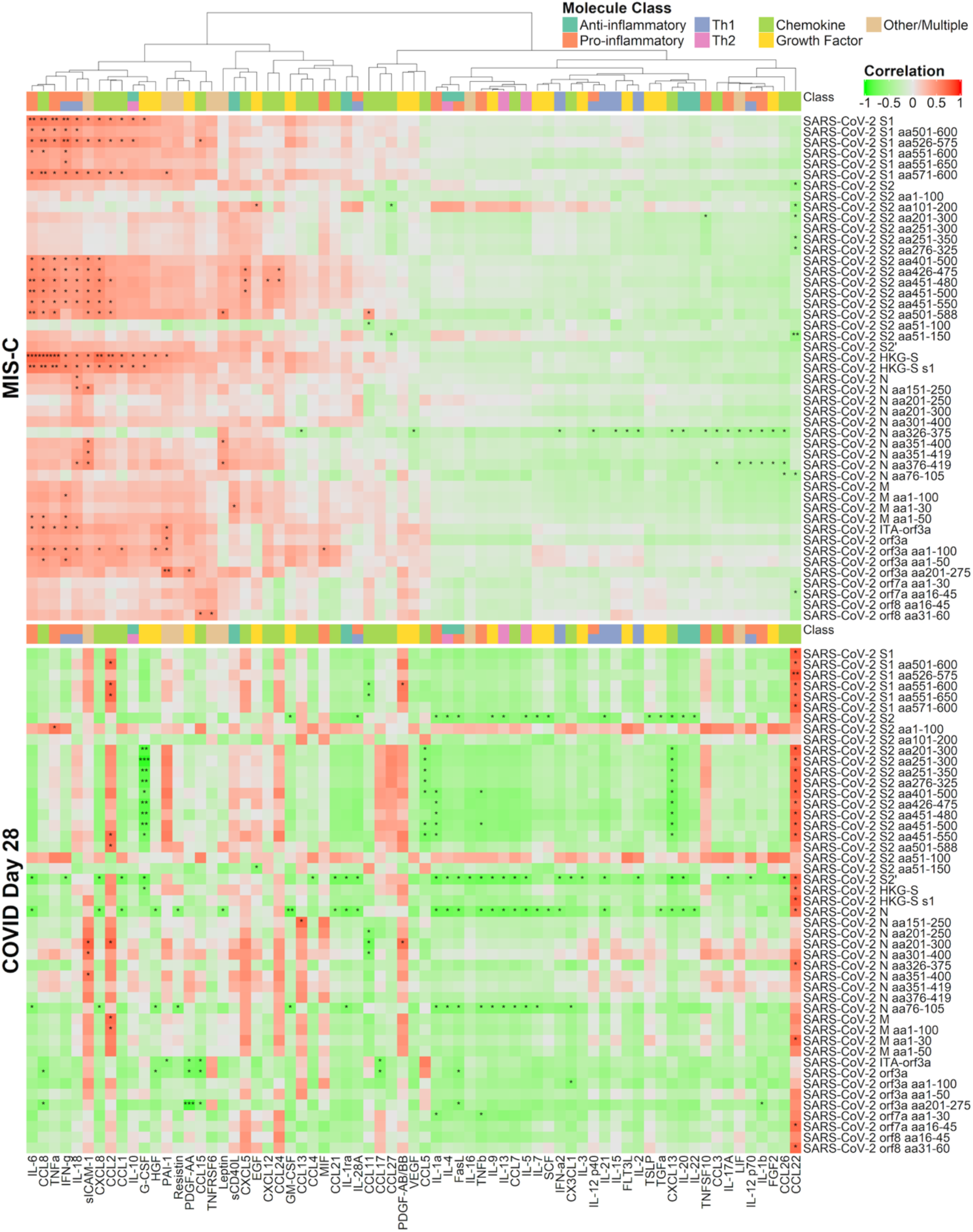
Correlation between IgG binding to SARS-CoV-2 proteins and fragments with selected cytokine levels. The heatmap shows the Pearson’s correlation coefficient between antibody binding levels and cytokine levels on a colorimetric scale. Significance of the correlations are shown by asterisks (*p<0.05, **p<0.005, ***p<0.0005). Plots are separated by patient group; MIS-C (top) and convalescent COVID-19, 28 days post-onset of symptoms (bottom). Antigens displayed correspond to proteins and fragments produced *in vitro* that were seropositive (≥ 1.0 normalized log_2_ signal intensity) in at least 20% of the study population. Cytokines were included if at least one significant association with the selected antibody responses was observed in either group of children. Cytokine classes are annotated at the top of the heatmap. The dendrogram at the top of the MIS-C group heatmap represents hierarchical clustering of the correlations.

## Discussion

The first human coronavirus (CoV) infections were described in the 1960s (Mulabbi et al., 2021). CoV infection of humans may result in either a simple “cold” or as seen with more recent variants, a life-threatening pandemic. Adding to the diversity of presentation and morbidity are the well-recognized differences in immune status and response based on age, portal of entry, underlying metabolic or immune deficiencies, previous exposures, and pathogenicity of the virus strain itself (Chou et al., 2022; Alefishat et al., 2022; Pierce et al., 2020; Tanner and Alfieri, 2021).

Data presented in this paper begin to address the differences observed between children with COVID-19 respiratory infections on or near day 28 post onset of symptoms, versus patients presenting with MIS-C approximately day 28 following exposure to SARS-CoV-2 during the first wave of infection in 2020. Ongoing research is centered on these time points, in order to begin to understand the possible role of diverse viral antigens in the contrasting immune response seen in SARS-CoV-2 respiratory syndrome versus MIS-C. Comprehending the differences in immune processing and response may aid in optimization of vaccines and may help develop targeted therapy for individuals at greater risk of severe respiratory infections or MIS-C associated with SARS-CoV-2 variants.

Our results demonstrate the association of humoral immunity to certain SARS-CoV-2 sequences with elevated cytokine and chemokine profiles in MIS-C versus day 28 following acute respiratory infection. The highly significant pattern of inflammatory cytokines which differentiated MIS-C from convalescent pediatric COVID-19 respiratory infection in our patients is similar to results reported elsewhere (Lapp et al., 2022; Conway et al., 2021; de Cevins et al 2021, Gurlevik et al., 2022; Lee et al., 2023). Our results are also in agreement with a similar study of antibody responses in MIS-C compared to pediatric COVID-19 (Thiriard et al, 2023), however, we investigated the IgG response to many more viral antigens, including structural and non-structural viral components and fragments of these proteins and correlated them with cytokine responses. It is clear that a combination of viral pathogenicity in balance with host immune response determines disease outcome; our approach now opens the way to study both, in patients across the continuum of age.

### Antigenicity

The lack of a sustained immune response to SARS-CoV-2 antigens has helped to promote the emergence of variants over time and has challenged researchers to come up with a more lasting vaccine. Even so, as in our studies, there are some antigens which dominate, and may play a role in triggering one pathway over another in the immune system (Jordan, 2021; Primorac et al., 2022). The striking, but steady decline in the incidence of MIS-C with each wave of variants over the past two years, may indicate that new viral epitopes are less likely than previous homologs to trigger the delayed and complex repertoire of humoral and cellular responses we now recognize in children 4-6 weeks after each spike in SARS-CoV-2 (Holm et al., 2022; Levy et al., 2022; Yuen et al., 2020). Alternatively, the drop in MIS-C cases over time may have resulted from the accumulation of pediatric SARS-CoV-2 infections during this period, which provided prior immunity that blocked the development of MIS-C. Our results largely agree with and add to previous studies of the immune profile of MIS-C (Anderson et al., 2021); (Lapp et al., 2022); (Ramaswamy et al., 2021; Thiriard et al, 2023: Gruber et al., 2020; Consiglio et al., 2020). Our study, however, used more MIS-C patients than all but one of these studies and investigated more cytokines and more coronavirus proteins than any previous study. Moreover, we were the first study to investigate antibody binding to fragments of SARS-CoV-2 proteins other than the RBD.

## Limitations

The limitations of this study include differences in the age, sex, BMI, time since infection and treatment in the two groups of patients we compared. We analyzed the effects of age, sex and BMI on IgG reactivity; only age showed significant effects. We therefore analyzed the contribution of age to IgG reactivity explicitly. We also analyzed the effects of age, sex and BMI on cytokine secretion and found a significant effect only for BMI on leptin levels in serum. This was significantly lower in MIS-C patients compared to 28-day convalescent pediatric COVID-19 patients primarily due to their lower BMI. We were not able to control precisely for time since infection or symptom onset since the MIS-C patients we studied initially had mild asymptomatic infections with SARS-CoV-2 that were not documented by PCR or antigen test. MIS-C patients were compared to convalescent pediatric COVID-19 patients 28 days post onset of symptoms since this time point best matched the known timing of MIS-C (Miller et al., 2022). Medical treatment was necessarily different for MIS-C patients compared to convalescent pediatric COVID-19 patients, but MIS-C patient samples were obtained prior to treatment and treatment had ceased at least two weeks earlier for the convalescent group, so it is unlikely that this had a significant effect. In our control patients, evidence of antibody reactivity from infused convalescent plasma was detected 24 hours post infusion, but was greatly reduced at twenty-one days post infusion (Figure S2).

## Conclusion

Our data show significant differences in coronavirus antibody responses in MIS-C patients compared to convalescent pediatric COVID-19 patients. MIS-C patients had lower levels of antibodies to endemic human coronaviruses, perhaps due to their younger age and fewer exposures to these viruses. Moreover, the MIS-C patients we studied had lower levels of antibodies reactive with several domains of the SARS-CoV-2 N, S and M proteins. This lack of immunity may allow greater spread of SARS-CoV-2 infection thereby triggering the inflammatory state that is clinically defined as MIS-C and characterized by the increased expression of inflammatory cytokines and chemokines that we noted. Our study included fine structure analysis of antibody binding to SARS-CoV-2 proteins, analysis of binding to HCoV proteins and extensive cytokine assays to characterize the differences in immunity between MIS-C patients and convalescent pediatric COVID-19 patients at or near the same time post infection.

## Acknowledgements

We thank Dr. Alvaro Galvis for clinical care of patients and for facilitating this study.

